# Seeding Activity of Skin Misfolded Tau as a Novel Biomarker for Tauopathies

**DOI:** 10.1101/2023.09.07.556724

**Authors:** Zerui Wang, Ling Wu, Maria Gerasimenko, Tricia Gilliland, Zahid Syed Ali Shah, Steven A. Gunzler, Vincenzo Donadio, Rocco Liguori, Bin Xu, Wen-Quan Zou

## Abstract

Tauopathies are a group of age-related neurodegenerative diseases with a molecular hallmark of the prion-like propagation and accumulation of pathologically phosphorylated tau protein in the brain. They include Alzheimer’s disease (AD), progressive supranuclear palsy (PSP), corticobasal degeneration (CBD), and Pick’s disease (PiD). Currently, in the peripheral tissues and body fluids there are no reliable diagnostic biomarkers available that are able to directly reflect the capability of propagation and spreading of the misfolded tau aggregates. Here, we revealed significantly increased amounts of phosphorylated tau in the skin of AD patients compared to those in other tauopathies and normal controls. Moreover, the seed-amplification assay (SAA) by the ultrasensitive real-time quaking-induced conversion (RT-QuIC) displayed that the prion-like seeding activity of pathological tau in the skin of cadavers with neuropathologically confirmed tauopathies including AD, PSP, CBD, PiD was dramatically higher than that in normal controls, yielding 75-80% sensitivity and 95-100% specificity, respectively, depending on different tau substrates used. The increased tau-seeding activity was also observed in biopsy skin samples from living AD and PSP patients. Moreover, analysis of the end products of skin-tau SAA confirmed that the increased seeding activity is accompanied with formation of tau aggregates that are of different physicochemical properties determined by the different tau-substrates used. Our study provides proof-of-concept that the skin tau-SAA can differentiate tauopathies from normal controls, suggesting that the seeding activity of the skin misfolded tau can serve as an accurate diagnostic biomarker of tauopathies.

## Introduction

The deposition of disease-associated tau aggregates in the brain is a pathological hallmark of tauopathies including Alzheimer’s disease (AD), Pick’s disease (PiD), progressive supranuclear palsy (PSP), and corticobasal degeneration (CBD) [23]. They share a common pathogenesis involving seeding and propagation of misfolded and highly phosphorylated isoforms of the cellular tau protein in the brain, a feature characteristic of infectious prion protein (PrP^Sc^ or prion) in prion disease (PrD) [42]. Human brain expresses six tau isoforms, resulting from three or four microtubule-binding repeats (3R or 4R tau) and 0-2 N-terminal inserts (0N, 1N, or 2N tau) [17]. It has been noticed that different tauopathies can have different composition of these tau isoforms. For instance, neuronal tau inclusions of the AD brain are composed of both 3R and 4R isoforms. In contrast, neuronal deposits of PiD primarily contains 3R isoforms, whereas PSP and CBD are characterized by the accumulation of 4R tau assembly in the brain. Different composition and morphology of tau isoforms among tauopathies are proposed to be associated with the existence of distinct strains of neurotoxic tau aggregate conformers [28]. A definite diagnosis of tauopathies relies on the availability of the brain tissue by biopsy or at autopsy for detection of tau-pathology and phosphorylated tau in the brain, which is highly invasive or too late. Recent advances in brain molecular imaging and highly sensitive immunoassays of phosphorylated and total tau (p-tau and t-tau) in plasma and cerebrospinal fluid (CSF) have made early and reliable AD diagnosis in living patients possible [56]. However, CSF sampling requires highly invasive lumbar puncture, while brain imaging is expensive and/or involves radioactivity. Newly developed single molecular immunoassay (Simoa) is highly sensitive and specificl in detection of the blood biomarkers, but Simoa is unaffordable to the majority of patients, especially those in the developing countries.

The ultrasensitive technology termed seed-amplification assay (SAA) of misfolded proteins by either real time quaking-induced conversion (RT-QuIC) or protein misfolding cyclic amplification assay (PMCA) has made it possible to identify new biomarkers in easily accessible specimens for early diagnosis and assessment of disease progression [36, 41, 57]. It has been widely used in the detection not only of prions but also other prion-like misfolded proteins such as αSyn and tau in body fluids and peripheral tissues of PrD and PD by detecting the seeding activity (SA) of misfolded proteins [4, 14, 18, 39, 40, 47, 50]. It has been reported by several labs that RT-QuIC can also detect tau-SA in the brain tissues from cadavers with tauopathies [25, 32, 45, 46, 53]. However, to date there has been no report of the application of tau-SAA to body fluids and peripheral tissues of AD patients although there was a report about tau RT-QuIC detection of CSF from PSP and CBD [46].

In the current study, we determine tau-SA of autopsy scalp skin samples from a cohort of neuropathologically confirmed cases comprising AD, PSP, CBD, PiD, and NC, using RT-QuIC with truncated truncated human tau constructs fragments as substrates. Our skin tau RT-QuIC reveals that misfolded tau from AD and other tauopathies like PSP and CBD, but not from normal controls and PiD, is selectively seeded by the 4RCF (equivalent to the human 4R tau fragment, K18CFh) substrate [25, 53] achieving a sensitivity of 80.5% and a specificity of 95.4%. With the 3RCF (equivalent to the human 3R tau fragment, K19CFh) substrate [45, 53], we display a sensitivity of 77% and a specificity of 92%. We also examine biopsied skin samples, including AD, PSP, and controls, and find high diagnostic efficacies for AD and controls. Moreover, we find increased skin tau-SAA in PD and PSP but not in PiD and normal controls. Our investigation constitutes the first application of SAA methodology to both post-mortem and biopsied cutaneous specimens and reveals potential of skin tau-SAA as a novel biomarker for diagnosis of tauopathies.

## Materials and Methods

### Ethical statement

All procedures and protocols were monitored and approved by the Institutional Review Boards (IRBs) of University Hospitals Cleveland Medical Center, Banner Sun Health Research Institute, and IRCCS Institute of Neurological Sciences of Bologna. Written informed consent was obtained from all living subjects undergoing skin biopsy or from family members for skin autopsy. For post-mortem sample collection, we obtained the specimens with respect to the wishes of the deceased and their family, following all legal and ethical guidelines. For skin biopsy procedures, all participants provided their informed consent prior to their inclusion in the study.

### Reagents and antibodies

Proteinase K (PK) and guanidine hydrochloride (GdnHCl) were purchased from Sigma Chemical Co. (St. Louis, MO, USA). Reagents for enhanced chemiluminescence (ECL Plus) were from Amersham Pharmacia Biotech, Inc. (Piscataway, NJ). Anti-tau mouse monoclonal antibodies RD3 and RD4 (Sigma-Aldrich) against human tau repeating region and sheep anti- mouse (SVM) IgG conjugated with horseradish peroxidase as a secondary antibody (AC111P, CHEMICON International, Inc, Burlington, MD, USA) were used. Antibodies against Phospho-Tau (Thr231) and phospho-Tau (Ser396) were purchased from Cell Signaling Technology (Danvers, MA, USA).

### Source of skin samples

A total of 135 autopsy scalp skin samples from AD (n=46), PSP (n=33), CBD (n=5), PiD (n=6) and non-neurodegenerative controls (NNCs, n=46) were collected and examined. These samples were obtained from the Arizona Study of Aging and Neurodegenerative Disorders (ASAND)/Brain and Body Donation Program at Banner Sun Health Research Institute through the Biomarkers across Neurodegenerative Diseases Research Grant 2019 (BAND 3) study. The diagnoses of these cases were confirmed via neuropathological examination of autopsied brain tissues at the ASAND. Biopsied skin samples from C7 paravertebral site (5 cm from the midline) of clinically diagnosed AD (n=16), PSP (n=8) and normal controls (n=10) were from the Bellaria Hospital, Bologna, Italy, and the University Hospitals Cleveland Medical Center, Cleveland, Ohio, USA (see neuropathological and clinical information in Tables 1 and 2).

**Table 1.**
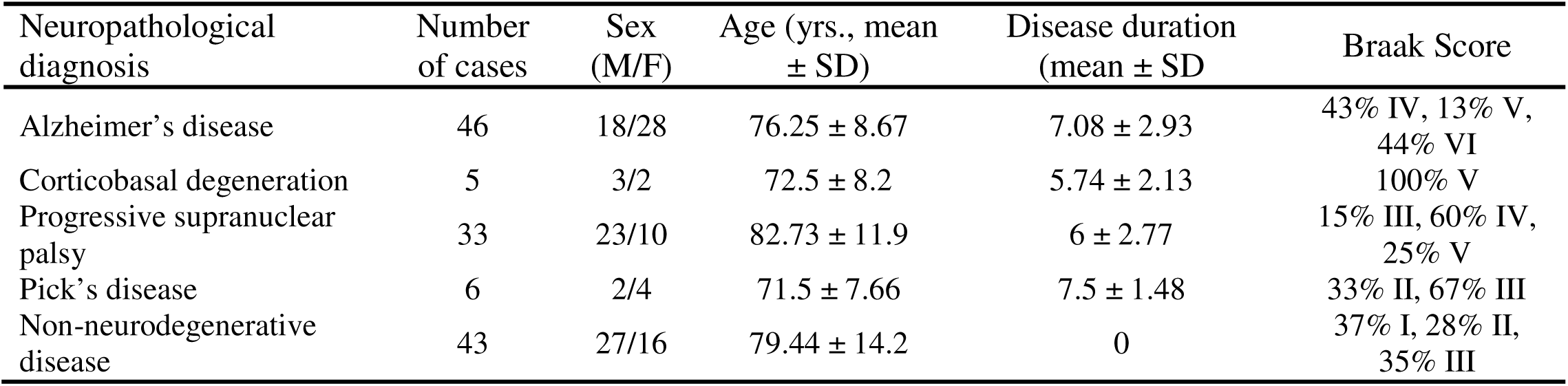
Demographic and neuropathological features of autopsied cases in different groups.

| Neuropathological diagnosis | Number of cases | Sex (M/F) | Age (yrs., mean ± SD) | Disease duration (mean ± SD) | Braak Score |
| --- | --- | --- | --- | --- | --- |
| Alzheimer's disease | 46 | 18/28 | 76.25 ± 8.67 | 7.08 ± 2.93 | 43% IV, 13% V, 44% VI |
| Corticobasal degeneration | 5 | 3/2 | 72.5 ± 8.2 | 5.74 ± 2.13 | 100% V |
| Progressive supranuclear palsy | 33 | 23/10 | 82.73 ± 11.9 | 6 ± 2.77 | 15% III, 60% IV, 25% V |
| Pick's disease | 6 | 2/4 | 71.5 ± 7.66 | 7.5 ± 1.48 | 33% II, 67% III |
| Non-neurodegenerative disease | 43 | 27/16 | 79.44 ± 14.2 | 0 | 37% I, 28% II, 35% III |

**Table 2.** Demographic and clinical features of biopsied cases in different groups.

| Clinical diagnosis | Number of cases | Sex (M/F) | Age (yrs., mean ± SD) | Disease duration (yrs., mean ± SD) | MMSE (mean ± SD) | Hoehn & Yahr stage (mean ± SD) |
| --- | --- | --- | --- | --- | --- | --- |
| Alzheimer's disease | 16 | 6/10 | 70.13 ± 9.68 | 7.08 ± 2.93 | 19.35 ± 5.44 | 0 |
| Progressive supranuclear palsy | 8 | 5/3 | 75.71 ± 9.36 | 6 ± 2.77 | 25.67 ± 4.93 | 3.27 ± 1.44 |
| Non-neurodegenerative disease | 10 | 3/7 | 68.80 ± 10.83 | 0 | Not available | 0 |

### Plasmid constructs cloning

Expression vectors for all six full-length wild-types human tau isoforms were generously provided by Dr. George Bloom of the University of Virginia (originated from the late Dr. Lester “Skip” Binder and Dr. Nicolas Kanaan of Michigan State University) [52, 53]. 3RCF construct (three microtubule-binding repeats and cysteine-free construct containing C322S mutation) was first PCR-amplified of 3R repeats sequence from 2N3R tau plasmid and cloned into the same expression vector using Nde I and Xho I restriction sites, followed by site-directed mutagenesis at Cys322 site to Serine using QuikChange Site-directed mutagenesis kit (Agilent, Santa Clara, CA; Fig. 1). 4RCF construct (four microtubule-binding repeats and cysteine-free construct containing C291S and C322S mutations) was first PCR-amplified of 4R repeats sequence from 2N4R tau plasmid and cloned into the same expression vector using Nde I and Xho I restriction sites, followed by site-directed mutagenesis at Cys291 and Cys322 sites using QuikChange Site-directed mutagenesis kit (Fig. 1). All constructs were designed with a his_6_-tag at their carboxy-termini to facilitate protein purification and were verified by DNA sequencing.

**Fig. 1.**
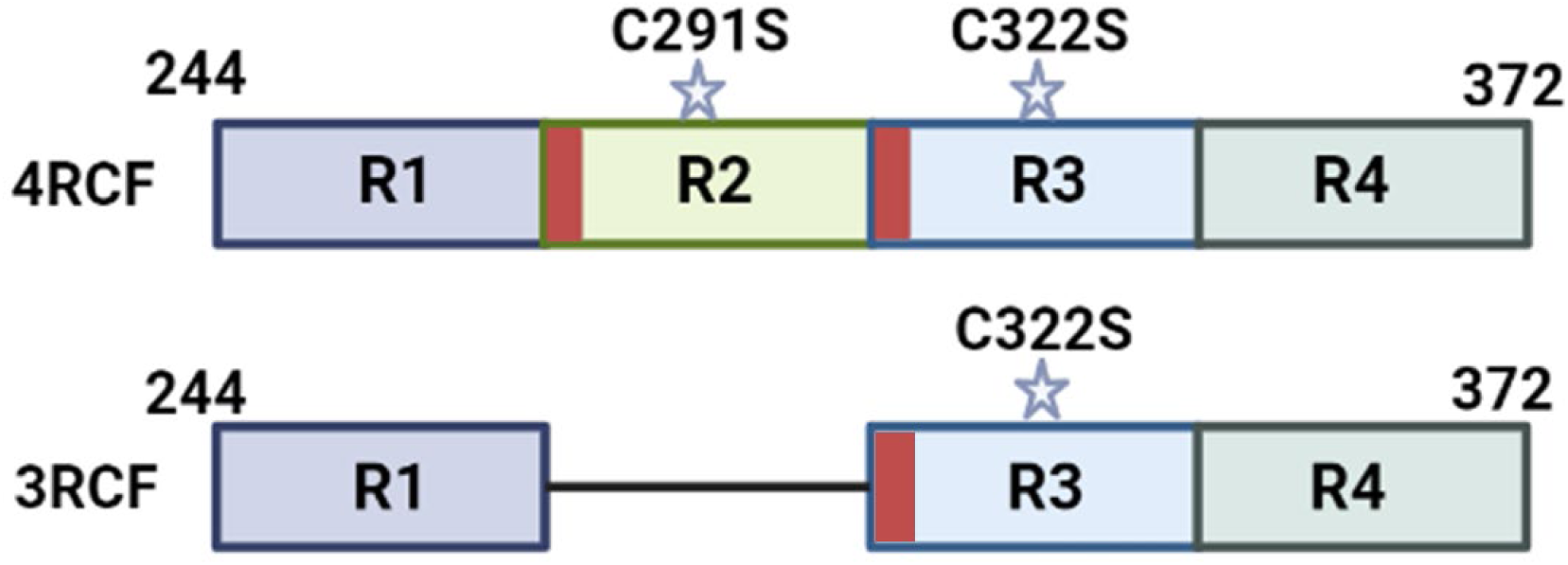
Schematic representation of truncated tau proteins. Diagram of recombinant 4RCF and 3RCF tau fragments with cysteine substitution for serine at residues 291 (C291S) and 322 (C322S). Both 4RCF and 3RCF start at residue 244 and ends at residue 372 but the latter lacks R2 domain.

### Engineered tau fragments 3RCF and 4RCF expression and purification

Recombinant 3RCF and 4RCF was prepared as previously described [53]. In brief, plasmids encoding human tau engineered constructs 3RCF and 4RCF were transformed into BL21-DE3 E. coli cells. Overnight starter cultures of BL21-DE3 E. coli cells transformed with recombinant tau plasmids were inoculated into multi-liter LB broth at 1:50 dilution and 100 mg/mL ampicillin. Cultures were incubated at 37°C, shaking until OD_600_ reached between 0.5-0.6. Tau expression was induced using 1 mM IPTG and continued to grow for an additional 4 hours. BL21-DE3 cells containing expressed tau were pelleted and resuspended in 50 mM NaH_2_PO_4_, pH 8.0 and 300 mM NaCl (sonication lysis buffer) at a concentration of 20 mL/L of culture preparation and sonicated at 60% power in ten 30-second intervals over 10 minutes. Cell lysates were centrifuged and supernatant containing the protein was applied to Ni-NTA column equilibrated with sonication lysis buffer. The columns were washed with 40-50 times of bed volumes of column buffer followed by washing buffer (50 mM NaH_2_PO_4_, pH 8, 300 mM NaCl, and 20 mM imidazole). Recombinant protein was then eluted using elution buffer (50 mM NaH_2_PO_4_, pH 8, 300 mM NaCl, and 200 mM imidazole). Fractions were tested for protein concentration using 5 μL of protein sample mixed with 10 μL Coomassie Protein Assay reagent (ThermoFisher Scientific). Pooled fractions were concentrated to 4 mL using 10 kDa molecular weight cut-off spin columns (Millipore) and filtered using 0.22 μm low-binding Durapore PVDF membrane filters (Millipore). 3RCF and 4RCF tau proteins were further purified by FPLC using size exclusion Superdex-75 and Superdex-200 columns (GE Healthcare) in 1 x PNE buffer (25 mM PIPES, 150 mM NaCl and 1 mM EDTA, pH 7.0). Final 3RCF and 4RCF proteins were over 90% purity as evaluated by SDS-PAGE. Protein concentrations were quantified by BCA protein assays (ThermoFisher Scientific).

### Skin tissue preparation

Skin samples of approximately 30-100 mg in weight and 3-5 mm x 3-5 mm in size, primarily contained epidermis and dermis were collected as previously and prepared described [49]. Briefly, skin tissues were homogenized at a 10% (w/v) concentration in a lysis buffer containing 2 mM CaCl_2_ and 0.25% (w/v) collagenase A (Roche) in Tris-Buffered Saline (TBS). The samples were incubated in a shaker at 37°C for 4 hours, shaking at 500 rpm, followed by homogenization using a Mini-BeadBeater (BioSpec, Laboratory Supply Network, Inc., Atkinson, NH, USA).

### RT-QuIC Analysis

The RT-QuIC assay was modified as previously described with a slight modification [25, 34, 45, 46, 53]. In brief, the reaction mix for skin tau was prepared with 10 mM HEPES, pH 7.4, 200 mM NaCl, 10 µM thioflavin T (ThT), and 10 µM either 4RCF or 3RCF tau substrate. In a 96-well plate (Nunc), 98 µL aliquots of the reaction mix were added to each well, followed by seeding with 2 µL of diluted skin homogenate (1:200 from 5% homogenate supernatant) in 10 mM HEPES, 1 x N2 supplement (Gibco), 1 x PBS and centrifuged at 5,000 g for 5 min at 4 °C. The plate was sealed with a plate sealer film (Nalgene Nunc International) and then incubated at 37 °C in a BMG FLUOstar Omega plate reader. The incubation involved cycles of 1 min of orbital shaking followed by a 15-min of rest for the specified duration. ThT fluorescence measurements from bottom read (450 ± 10 nm excitation and 480 ± 10 nm emission) were recorded every 45 min. Each sample dilution contained 4 replicate reactions. The average ThT fluorescence values per sample were calculated using data from all four replicate wells, regardless of whether they crossed the threshold defined by ROC. A sample was considered positive if at least 2 of 4 replicate wells exceeded this threshold.

To quantify tau-SA detected by RT-QuIC, end-point dilution titrations were employed to determine the estimates of the sample dilution that generated positive reactions in 50% of the replicate reactions as the 50% seeding dose or SD_50_ (usually 2 out of 4 replicates) [50].

### Conformational stability immunoassay

The conformational stability immunoassay of RT-QuIC end product was conducted as previously described with a minor modification [59]. Briefly, 20 µL aliquots of end product were mixed with 20 µL of GdnHCl stock solution, resulting in final GdnHCl concentrations ranging from 0 to 3.0 M. After incubating at room temperature for 1.5 hours, samples were precipitated with a 5-fold volume excess of pre-chilled methanol overnight at −20°C. Following centrifugation at 14,000 g for 30 minutes at 4°C, the pellets were resuspended in 20 µL of lysis buffer (10 mM Tris-HCl, 150 mM NaCl, 0.5% Nonidet P-40, 0.5% deoxycholate, 5 mM EDTA, pH 7.4). Each aliquot was digested with 10 µg/mL PK for 30 minutes at 37°C. The reaction was terminated with cOmplete protease inhibitor cocktail (CO-RO, Roche), and the samples were boiled in SDS loading buffer and loaded onto 15% Tris-HCl pre-cast gels (Bio-Rad) for Western blotting analysis.

### Velocity sedimentation in sucrose step gradients

RT-QuIC end products were mixed with 20% Sarcosyl to achieve a final concentration of 2% Sarcosyl as previously described [55]. Each sample was loaded onto 10-60% step sucrose gradients and centrifuged at 200,000 × g using an SW55 rotor for 1 hour at 4°C, with a minor modification to the described method [55]. After ultracentrifugation, the contents of the centrifuge tubes were sequentially removed from top to bottom, yielding 12 fractions that were subjected to Western blotting as detailed below.

### Western blotting

The samples prepared as described above were separated using 15% Tris-HCl Criterion pre-cast gels (Bio-Rad) in SDS-PAGE. Proteins from the gels were transferred onto Immobilon-P polyvinylidene fluoride (PVDF, Millipore) membranes for 90 minutes at 70 V. To probe with the anti-tau antibodies (RD3, RD4, pT231, or pS396), the membranes were incubated overnight at 4°C with a 1:1,000-1:4,000 dilution of the primary antibody. After incubation with a 1:4,000-1:5,000 dilution of horseradish peroxidase-conjugated sheep anti-mouse IgG, tau bands were visualized on Kodak film using ECL Plus as instructed by the manufacturer. Densitometric analysis was used to measure the intensity of tau protein bands, which were quantified with UN-SCAN-IT Graph Digitizer software (Silk Scientific, Inc., Orem, Utah).

### Filter-trap assay

The filter-trap assay was used to determine the RT-QuIC reaction mixtures with increased ThT fluorescence formed aggregates and to evaluate their sizes as described previously [38]. In brief, the end products of RT-QuIC were mixed with a 2% SDS buffer and incubated for the desired duration under suitable conditions to promote aggregation. Following incubation, the samples were filtered through a cellulose acetate membrane (Advantec MFS, Dublin, CA). After filtering, the membrane was rinsed with 0.1% SDS washing buffer to remove unbound proteins and subsequently blocked with 5% BSA for an hour. The membrane was then probed with RD3 and RD4 antibodies, followed by incubation with sheep anti-mouse secondary antibody. The proteins on the membrane were visualized using ECL Plus, and the resulting signal was captured using a chemiluminescent imaging system with X-ray/automatic film processor. Densitometric analysis was used to measure the intensity of tau protein dots for the quantitative analysis as mentioned above.

### Transmission electron microscopy analysis

Transmission electron microscopy (TEM) images were collected as previously described [53, 58]. Briefly, the skin tau-SAA end product samples at a concentration of 10 µM were maintained in a frozen state until ready for TEM analysis. Before imaging, 2 µL of the sample was applied to a 200 mesh formvar-carbon coated grid and allowed to sit for 5 minutes. The method of negative staining was used for imaging. The excess sample was gently removed using filter paper. A thorough examination of each grid was conducted to qualitatively assess the presence of oligomers or fibrils. Representative images were taken from 15–20 distinct locations on each grid. The TEM studies were conducted using a JEOL-1400 transmission electron microscope (JOEL United States, Inc., Peabody, MA) at an operating voltage of 120 kV.

### Statistical analysis

Experimental data were analyzed using Student’s t-test for comparing two groups. McNemar’s test was employed to assess marginal homogeneity and differences in agreement. For comparisons between PD versus CBD and PSP, where the sample size allowed, we conducted a paired area under the ROC curve (AUC) analysis to evaluate significant differences in AUC values. Tests adopted a two-sided type II error level of 0.05.

## Results

### Phosphorylated-tau is detectable in skin samples of patients with tauopathies

We first determined whether the pathologically phosphorylated tau is detectable in the skin samples of patients with tauopathies by Western blotting. Skin homogenates from cadavers with AD (n=5), PSP (n=4), CBD (n=4), PiD (n=4) and NC (n=4) were examined by western blotting with antibodies directed against phosphorylated tau (Anti-pT231 and Anti-pS396). P-tau231 and p-tau396 epitopes were previously identified from a systematic p-tau antibody screening in differential detection with AD and control brains [51]. Both the pThr231 (Fig. 2a) and pSer396 (Fig. 2b) antibodies revealed protein bands migrating at between approximately 25 and 110 kDa in all AD, 1 of 4 PSP, 2 of 4 CBD, and 1 of 4 PiD skin samples (Fig. 2). These bands may represent truncated and full-length monomers as well as oligomers of phosphorylated tau. In contrast, normal controls mainly displayed a band migrating at about 110 kDa. The levels of phosphorylated tau in the skin of AD patients were significantly elevated compared to those of other tauopathies and normal controls, as evidenced by the quantitative analysis of all phosphorylated tau bands in each lane on the pT231 blot with (Fig. 2c) and without (Fig 2d) the top band migrating at 110 kDa. Similarly, higher levels of phosphorylated tau were detected by p-tau396 in AD than those in other tauopathies for all bands from 25 kDa~110 kDa (Fig. 2e) and without (Fig. 2f) the top band at 110 kDa. To determine the specificity of the protein bands and exclude the possibility that the bands might be from the non-specific reaction of the secondary antibody, we probed these bands with the secondary antibody only in the absence of the primary antibody (Fig. S1). The absence of detectable bands following a long ECL exposure (90 minutes) suggested that the bands identified by phospho-tau antibodies were indeed specific. The results revealed that the levels of the skin phosphorylated tau were significantly greater in AD than in other tauopathies and controls.

**Fig. 2.**
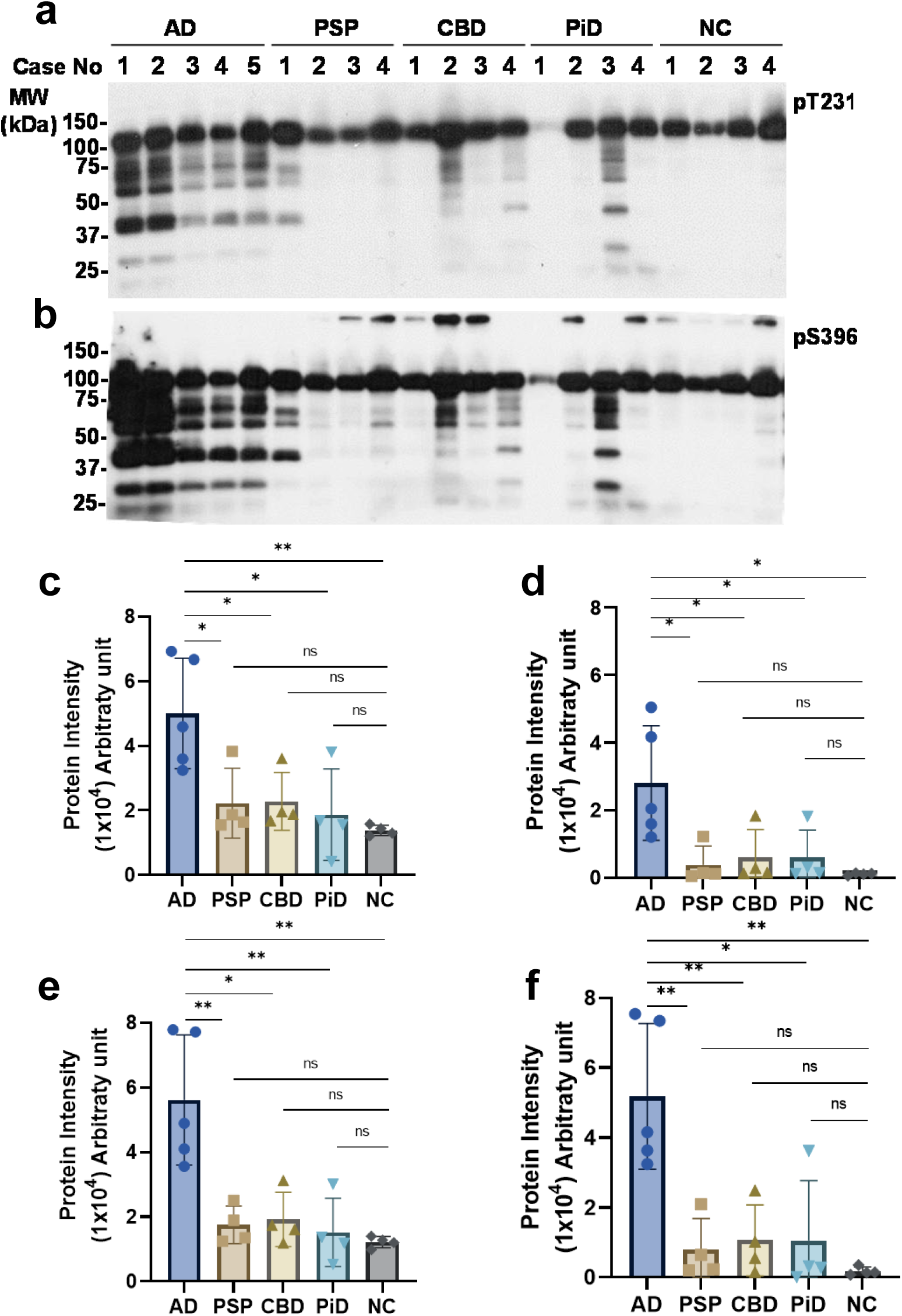
Western blotting of phosphorylated tau in the skin of cadavers with tauopathies. **a** Conventional western blotting of skin homogenates from AD (n=5), PSP (n=4), CBD (n=4), PiD (n=4) and NC (n=4), probed by anti-tau (pT231). **b** Skin homogenates from AD (n=5), PSP (n=4), CBD (n=4), PiD (n=4) and NC (n=4), immunoblotting with anti-tau (pS396). The specific bands migrated at 37~50 kDa were observed in all the AD cases but not in the non-neurodegenerative controls (NC). **c** Bar graph representing quantitative protein intensity of all bands in each lane on the blot probed with the pT231 antibody shown in (**a**). **d** Bar graph representing quantitative protein intensity of all bands in each lane except for the top band migrating at ~110 kDa on the blot probed with the pT231 antibody shown in (**a**). **e** Bar graph depicting quantitative protein intensity of all the bands in each lane on the blot probed with the pS396 antibody shown in (**b**). **f** Bar graph depicting quantitative protein intensity of all bands in each lane except for the top band migrating at ~110 kDa on the blot probed with the pS396 antibody shown in (**b**).

### Skin tau-seeding activity can specifically differentiate AD and non-AD tauopathies from normal controls and PiD cases using 4RCF as the substrate

In contrast to PrP in PrD and α-synuclein in PD, the tau molecule in the human brain exhibits 6 isoforms, of which there are 3 isoforms with 4 microtubule-binding repeats (4R tau) and 3 with three repeats (3R tau), resulting from alternative mRNA splicing. In order to find an appropriate tau substrate for setting up our seed-amplification assay of skin tau (sTau-SAA) with the ultrasensitive RT-QuIC technique, based on our recent finding with autopsy brain tissues from cadavers with AD and other non-AD tauopathies [53], we first examined 4 types of tau substrates including 2 full-length tau isoforms (2N3R/2N4R) and 2 truncated tau fragments consisting of 4RCF (cysteine-free, equivalent to K18CFh)/3RCF (equivalent to K19CFh) using autopsy skin samples from AD cadavers. 4RCF represents the core aggregation sequences of the 4R tau isoforms [51, 53]. Correspondingly, 3RCF fragment represents the core aggregation sequences of the 3R tau isoforms. The only difference between 4RCF and 3RCF is that 3RCF lacks R2 segment sequence. Compared to the full-length tau, the truncated tau substrates-based RT-QuIC revealed higher endpoint fluorescence readings (exceeding 100,000 RFU) and shorter lag phases (less than 20 hours) (Fig. S2). As a result, we used the two truncated tau fragments as the substrate for the following examinations in our study.

Using RT-QuIC with 4RCF or 3RCF truncated tau as the substrate, we analyzed autopsy skin samples from AD (n=42), CBD (n=4), PSP (n=33), PiD (n=6), and NC (n=43). 4RCF-based RT-QuIC of skin tau-seeding activity (sTau-SA) exhibited that CBD cases had the highest ThT fluorescence intensity, followed by AD, PSP, and PiD (Fig. 3a, b). We also compared lag phases of sTau-SA in each type of tauopathy, the time period between the beginning of the RT-QuIC measurement and the time point at which the curves reflecting the ThT fluorescence started to increase in their ThT fluorescence. Consistent with the endpoint ThT values, CBD displayed the shortest lag phase, while PiD was the longest (Fig. 3c). Our 4RCF-based RT-QuIC assay of the skin samples from AD and non-AD tauopathies yielded a sensitivity of 80.49% and a specificity of 95.35%. To quantitate the sTau-SA of each type of tauopathy, we conducted endpoint titration of RT-QuIC assay of each skin sample from different tauopathies. The x-axis represents SD_50_ that referred to the 50% seeding activity. PSP demonstrated the highest sTau-SA, followed by AD, while PiD skin samples showed the lowest tau-SA (Fig. 3d). AUC analysis revealed an area value of 0.88 based on the comparison between AD and normal controls and 0.79 based on the comparison between tauopathies and normal controls (Fig. 3e, f).

**Fig. 3.**
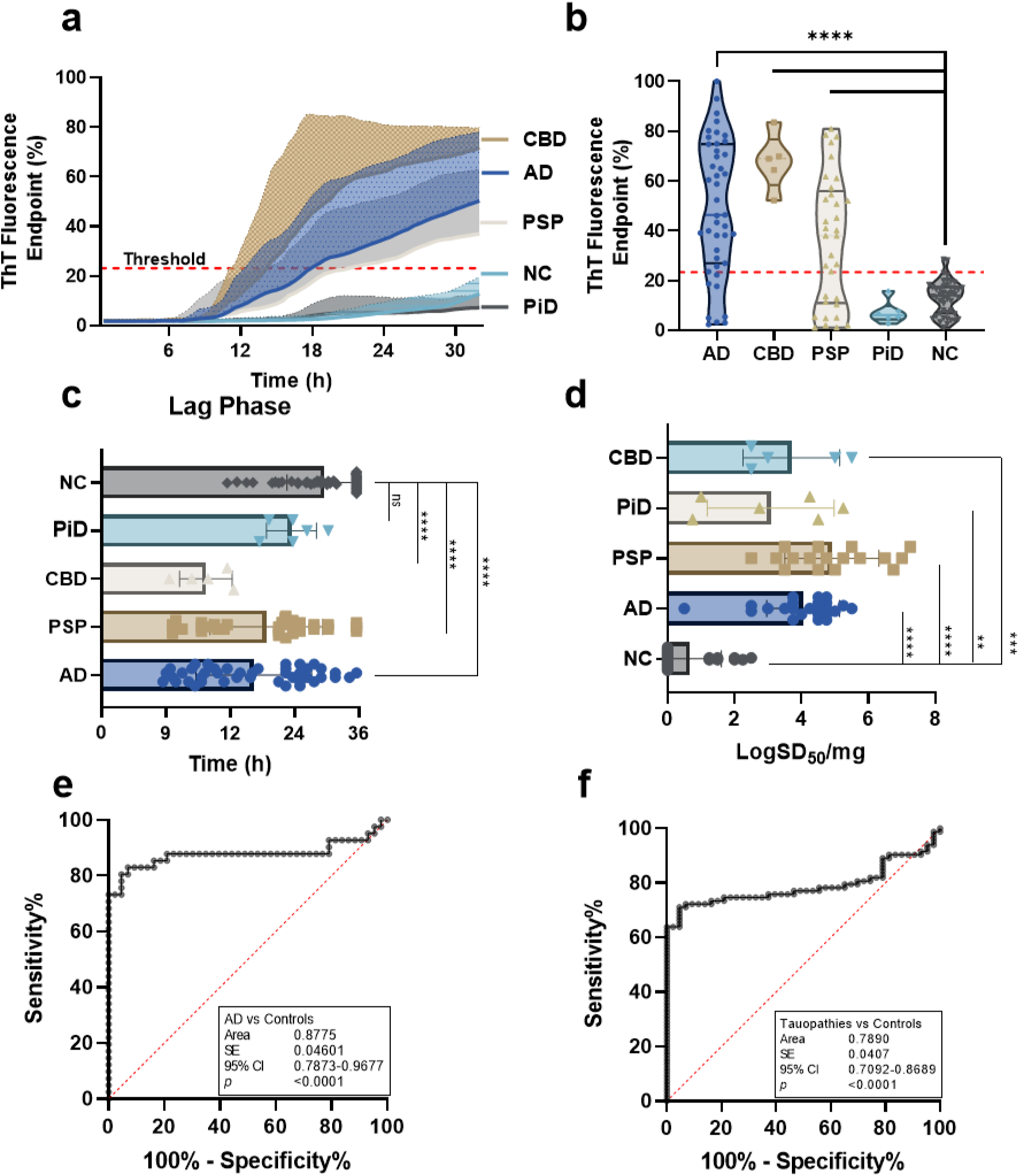
Tau-seeding activity of skin samples from patients with tauopathies using 4RCF-based RT-QuIC. **a** Kinetic curves displaying the mean and standard deviation (SD) of tau-SA over time of skin samples from CBD (n=5), AD (n=46), PSP (n=33), PiD (n=6) and NC (n=43). **b** Scatter plot illustrating the distribution of tau-SA across different tauopathies detected in panel (**a**). **c** Lag phase, defined as the initial period before a significant increase in the ThT fluorescence curves. **d** The end-point dilution analysis of quantitative tau-SA of skin samples from tauopathies. The half of maximal SA (SD_50_) was determined by Spearman-Kärber analyses and is shown as log SD_50_/mg skin tissue [18]. **e** Receiver operating characteristic (ROC) curve analysis comparing AD patients and control subjects, with an area under the curve (AUC) of 0.82. **f** ROC curve analysis comparing total tauopathies and control subjects, with an AUC of 0.79. ****: *p* < 0.0001; ***: *p* < 0.001; **: *p* < 0.01.

### The sTau-SA of AD and all tauopathies can be specifically differentiated from that of normal controls using 3RCF as the substrate

We next used 3RCF (K19)-based RT-QuIC assays to examine the same samples detected in 4RCF studies. The sTau-SA was significantly higher in tauopathies than in non-tauopathies, of which PiD was the highest, followed by AD, PSP, and CBD (Fig. 4a, b). We also compared their lag phases, which were inversely proportional to those of 4R tau: PiD exhibited the shortest lag phase, while CBD showed the longest lag phase (Fig. 4c). We also quantitated the skin 3RCF based tau-SA of each type of tauopathy by the endpoint titration of RT-QuIC assay of each skin sample from different tauopathies. PiD and PSP demonstrated the highest seeding dose while CBD had the lowest one (Fig. 4d). In addition, the 3RCF-based skin tau RT-QuIC assay generated a sensitivity of 75% and a specificity of 100%, which was similar to that of 4RCF-based RT-QuIC in general. The AUC values (0.85 vs 0.79) of 3RCF-based RT-QuIC assay of skin tau were similar to those detected with 4RCF-based RT-QuIC shown above (Fig. 4e, f). Notably, in contrast to the 4RCF-based sTau-SA, 3RCF-based sTau-SA from PiD was the highest, in addition to differentiating other tauopathies from the normal controls.

**Fig. 4.**
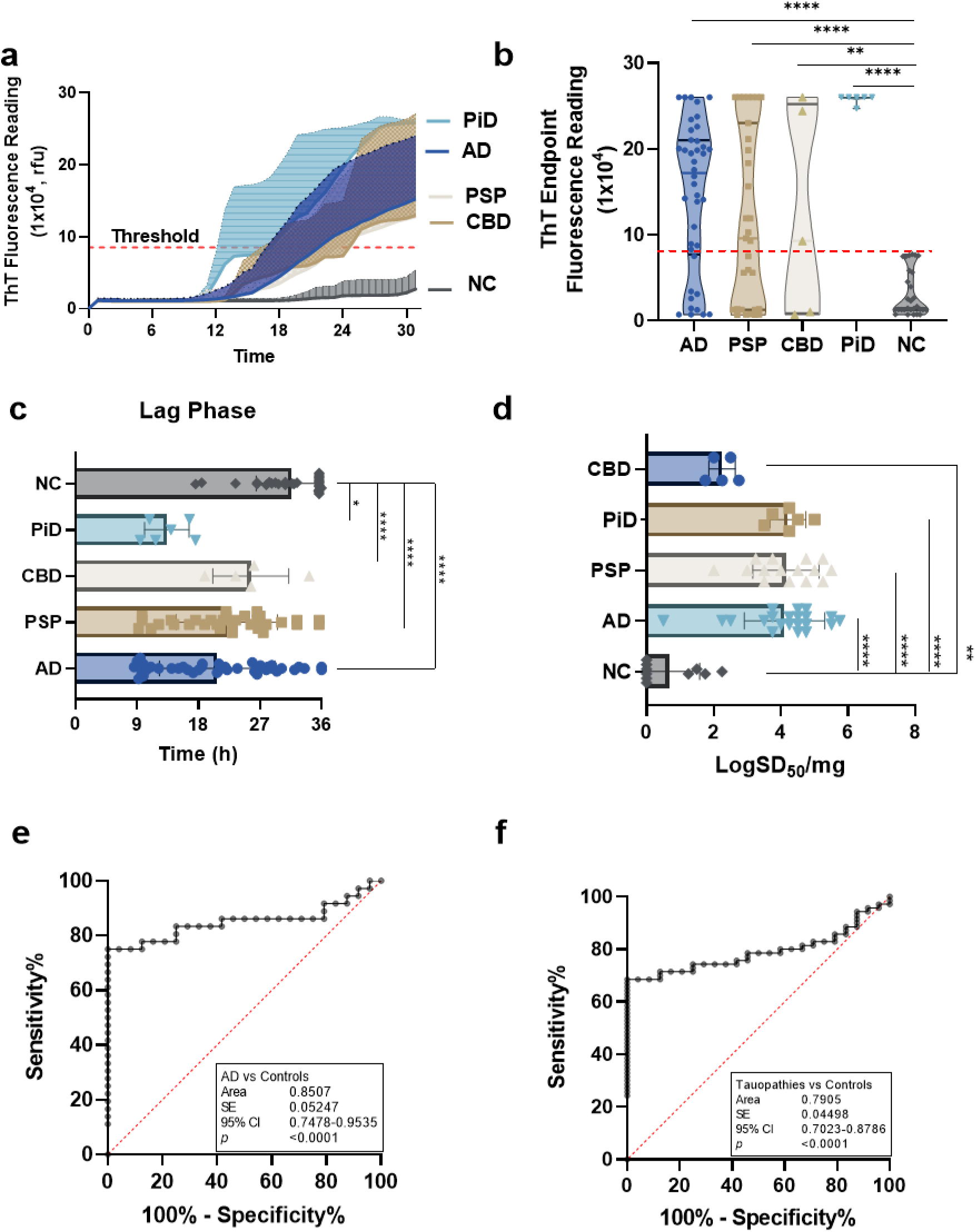
Tau-seeding activity of skin samples from patients with tauopathies using 3RCF-based RT-QuIC assay. **a** Kinetic curves displaying the mean and SD of tau-SA over time of skin samples from CBD (n=5), AD (n=46), PSP (n=33), PiD (n=6) and NC (n=43). **b** Scatter plot illustrating the distribution of tau-SA across different tauopathies. **c** Lag phase, same as above, as the initial delay before the ThT fluorescence curves begin to rise. **d** The end-point dilution analysis of quantitative tau-SA of skin samples from tauopathies. The half of maximal SA (SD_50_) determined by Spearman-Kärber analyses is shown as log SD_50_/mg skin tissue. **e** ROC curve analysis comparing AD patients and control subjects, with an AUC of 0.77. **f** ROC curve analysis comparing total tauopathies and control subjects, with an AUC of 0.72. *: *p* < 0.05; **: *p* < 0.01; ***: *p* < 0.001; ****: *p* < 0.0001.

### The sTau-SA is significantly higher in PD and dementia with Lewy bodies than in multiple system atrophy and normal controls but it is still lower than that in AD

Accumulation of tau aggregates in the brain has been observed in synucleinopathies including PD, dementia with Lewy bodies (DLB), and multiple system atrophy (MSA) [10, 19, 37, 44, 48]. Next, we further explored whether tau-SA can be detected in the skin of synucleinopathies by our sTau-SAA. Autopsy skin samples from AD (n=21), PD (n=10), MSA (n=6), DLB (n=6), and NC (n=17) were examined by 4RCF-based tau-SAA. The ThT endpoint fluorescence intensity of sTau-SAA was dramatically higher in AD than in synucleinopathies, whereas the skin-tau fluorescence intensity was significantly higher in synucleinopathies except for MSA than in the control group (Fig. 5), consistent with the previous observations that some of cases with synucleinopathies can have tau-pathology [10, 19, 37, 44, 48].

**Fig. 5.**
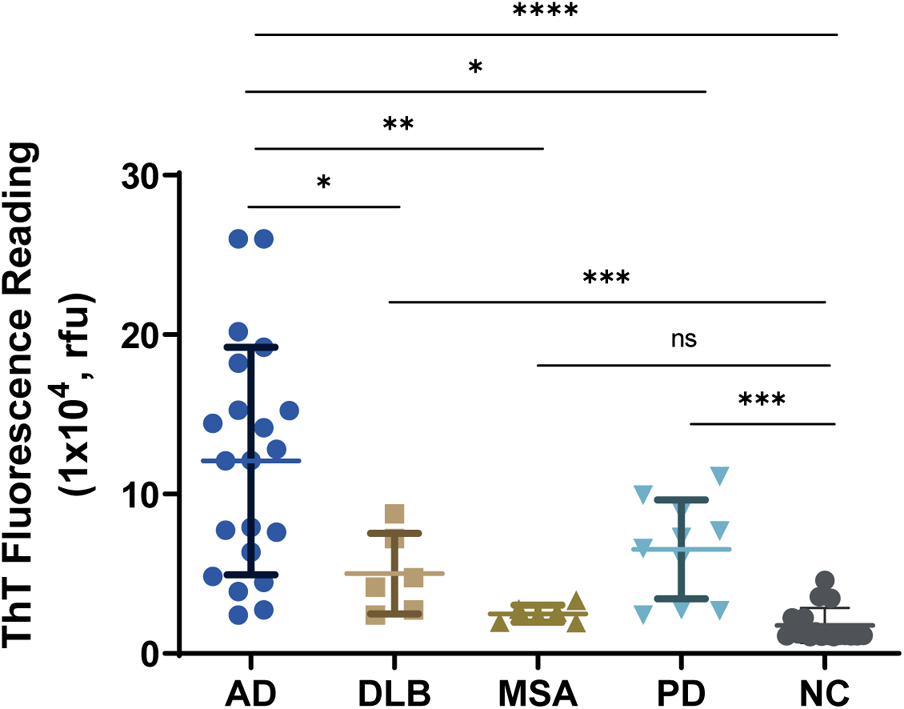
Tau-SAA of skin samples from patients with AD, Synucleinopathies and NC using 4RCF-based RT-QuIC assay. Scatter plot illustrating the distribution of tau-SA at the endpoint fluorescence readings across skin samples from 21 cases with AD, different synucleinopathies including 6 cases with DLB, 6 with MSA and 10 with PD as well as 17 NCs. *: *p* < 0.05, **: *p* <0.01, ***: *p* < 0.001; ****: *p* < 0.0001.

### Biopsy sTau-SA is significantly higher in tauopathies than in normal controls

We then used the above 4RCF- or 3RCF-based SAA (4RCF- or 3RCF-SAA) to examine biopsied skin samples from AD (n=16), PSP (n=8) and NCs (n=10). Skin 4RCF-SA was significantly higher in AD than in normal controls [83091 ± 57912 (mean ± SD) vs 20490 ± 9307, *p* = 0.0026 < 0.005]; skin 4RCF-SA was also significantly greater in PSP than in normal controls (73360 ± 49625 vs 20490 ± 9307, *p* = 0.0043 < 0.005) (Fig. 6a). Similar to skin 4RCF-SA, 3RCF-SA was significantly higher in AD than in NCs [97511 ± 54115 vs 30090 ± 13657, *p* = 0.0008 < 0.001] and greater in PSP than in NCs (60449 ± 20492 vs 30090 ± 13657, *p* = 0.0017 < 0.005) (Fig. 6b). There were no significant differences in sTau-SA between AD and PSP cases with both substrates (Fig. 6). This observation implied the potential for sTau-SA to serve as an antemortem diagnostic biomarker to differentiate tauopathies from normal controls.

**Fig. 6.**
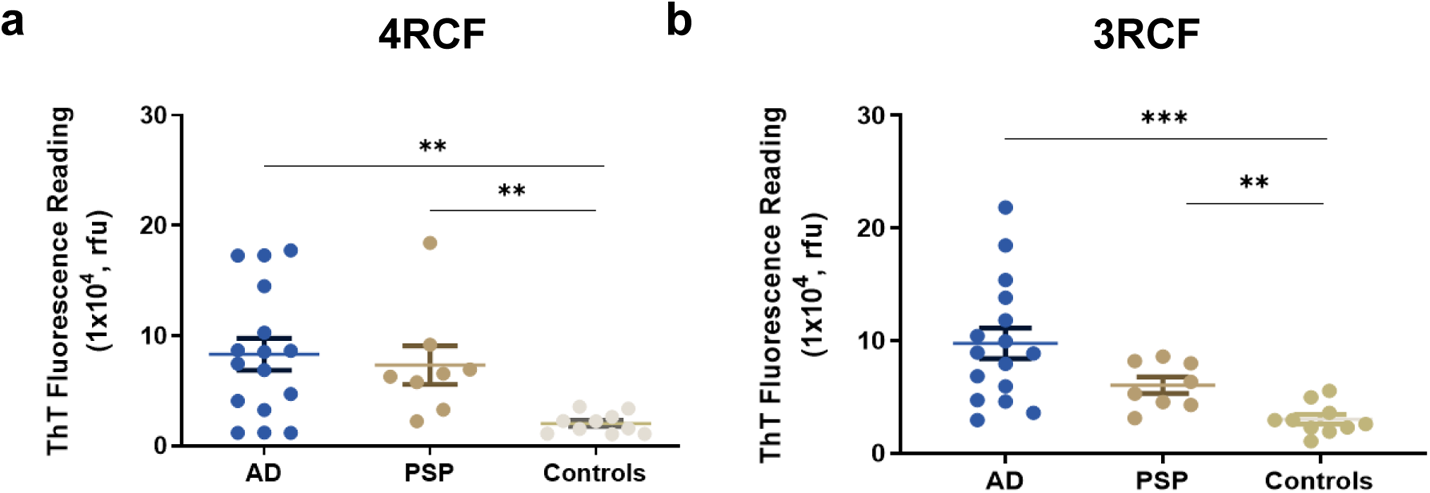
Examination of tau-SA in biopsied skin samples from patients with AD and PSP using either 3RCF- or 4RCF-based RT-QuIC assay. The scatter plot displays the endpoint ThT fluorescence intensity for AD (n=16), PSP (n=8), and normal control samples (n=10). **: *p* < 0.01; ***: *p* < 0.001.

### ThT fluorescence levels of sTau-SAA end-point correlate with dot-blot intensity of captured tau aggregate of the RT-QuIC end products

To determine whether ThT fluorescence levels reflecting sTau-SA represent the formation of skin tau-seeded aggregates, we correlated the end-point ThT fluorescence levels with the dot-blot intensity of tau aggregates captured by a filter-trap assay (FTA) (Fig. 7). After obtaining the end point ThT fluorescence levels of sTau-SA of 4 cases each from AD, PSP, CBD, PiD and NC with 4RCF or 3RCF as the substrate (Fig. 7a, b), we then ran FTA with their corresponding end products, followed by probing the dot-blots with anti-4R tau antibody (RD4) (Fig. 7c) and anti-3R antibody (RD3) (Fig. 7d). The semiquantitative densitometric scanning of protein dot intensity on the dot-blots revealed that similar to ThT fluorescence levels, the intensity of tau aggregates captured by FTA on the blots was significantly higher in tauopathies than in normal controls by both RD4 (Fig. 7e) and RD3 antibodies (Fig. 7f). Correlation analyses demonstrated that the intensity of the trapped aggregates from the end products correlated positively with the ThT fluorescence levels (r = 0.86 for 4R, r = 0.68 for 3R) (Fig. 7g, h).

**Fig. 7.**
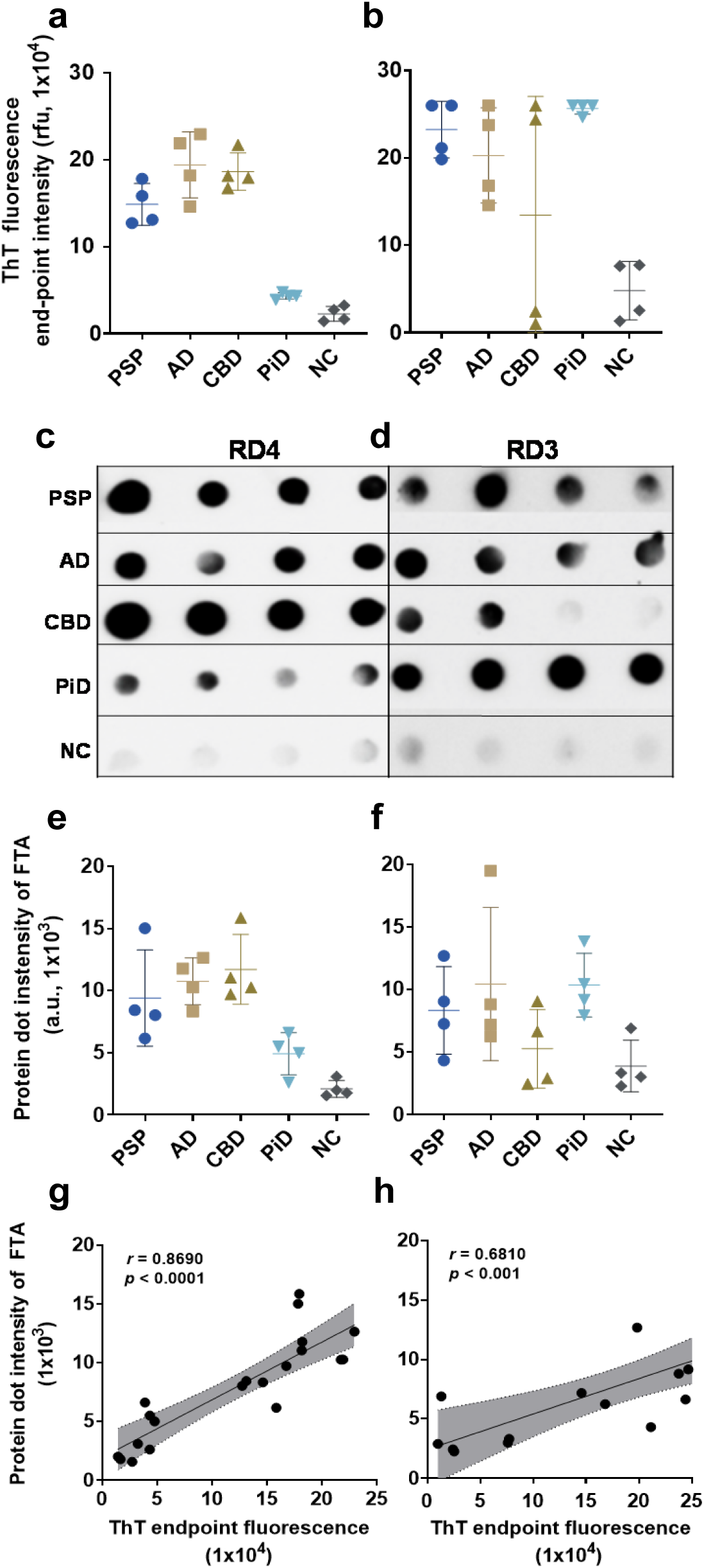
Characterization of RT-QuIC end products of skin tau from tauopathies using Filter-Trap Assay (FTA) probed with anti-3R (RD3) and anti-4R (RD4) tau antibodies. **a** and **b** Scatter plot of ThT fluorescence values skin-tau RT-QuIC end products of selected samples from PSP (n=4), AD (n=4), CBD (n=4), PiD (n=4) and NC (n=4), with (**a**) 4RCF and (**b**) 3RCF substrates. **c** and **d** FTA assay of 3RCF-/4RCF-based RT-QuIC end products of skin samples from different tauopathies including PSP, AD, CBD, PiD and normal controls (4 cases from each group) probed with RD4 (**c**) or RD3 (**d**) antibodies. **e**-**f** Densitometric quantification of density of FTA-dot blotting with 4RCF- (**e**) and 3RCF(**f**) - based RT-QuIC end products from panel **c** and **d**. **g**-**h** Correlation analysis between FTA-trapped protein dot intensity and skin tau-SA of 4RCF-/3RCF-based RT-QuIC end products.

### Transmission electron microscopy of 4RCF- or 3RCF-based RT-QuIC end products shows large oligomers but not mature fibrils

Although our FTA apparently was able to detect captured tau aggregates, transmission electron microscopy (TEM) detected only oligomer-like structures in 3RCF and 4RCF positive SAA end products (Fig. S3 a, b). In contrast, obviously matured fibrils of the end product of ɑSyn RT-QuIC were detectable by TEM (Fig. S3 c, d).

### The end products of 4RCF- and 3RCF-SAA exhibit different patterns of resistance to proteinase K digestion

The pattern of protein aggregates from the skin tau RT-QuIC end products to proteinase K (PK) digestion has been widely believed to reflect the conformational properties of misfolded proteins examined. Since the small fragments of 3RCF tau lower than 7 kDa are more difficult to digest by PK, we next performed the titration of varied PK concentrations ranging from 0, 1.25 µg/mL, 2.5 µg/mL, 3.75 µg/mL, 5 µg/mL, 7.5 µg/mL, to 10 µg/mL for the 4R (Fig. 8a) and 0, 1.25 µg/mL, 5 µg/mL, 10 µg/mL, 12.5 µg/mL and 25 µg/ml for the 3R tau (Fig 8b) SAA end products of skin tau from AD and non-AD subjects. In the end products treated without PK of 4RCF-based RT-QuIC of skin tau from non-AD, three protein bands were detected by the RD4 tau antibody, migrating at approximately 25-27 kDa, 12-14 kDa, and 7-10 kDa (Fig. 8a). The short exposure of our blots showed that the 7-10 kDa bands actually consisted of 7 kDa and 10 kDa proteins. As a result, the above four bands could represent the trimer and dimer of a 7 kDa band as well as the monomers of a full-length 4RCF (~12 kDa) and a truncated 4RCF (~7-10 kDa), respectively. Of them, the two lower monomeric bands were predominant, whereas the top dimeric and trimeric tau bands were underrepresented, accounting for less than 1-2% of total tau (Fig. 8a, c). In contrast, the end product from AD skin samples without PK-treatment exhibited an additional band migrating at approximately 48-50 kDa in addition to the four bands found in the end product of non-AD skin described above. This band could be oligomers of full-length or truncated tau molecules. In addition, the intensity of truncated trimer and dimer of tau migrating at 24-27 kDa was significantly increased compared to that of non-AD end product.

**Fig. 8.**
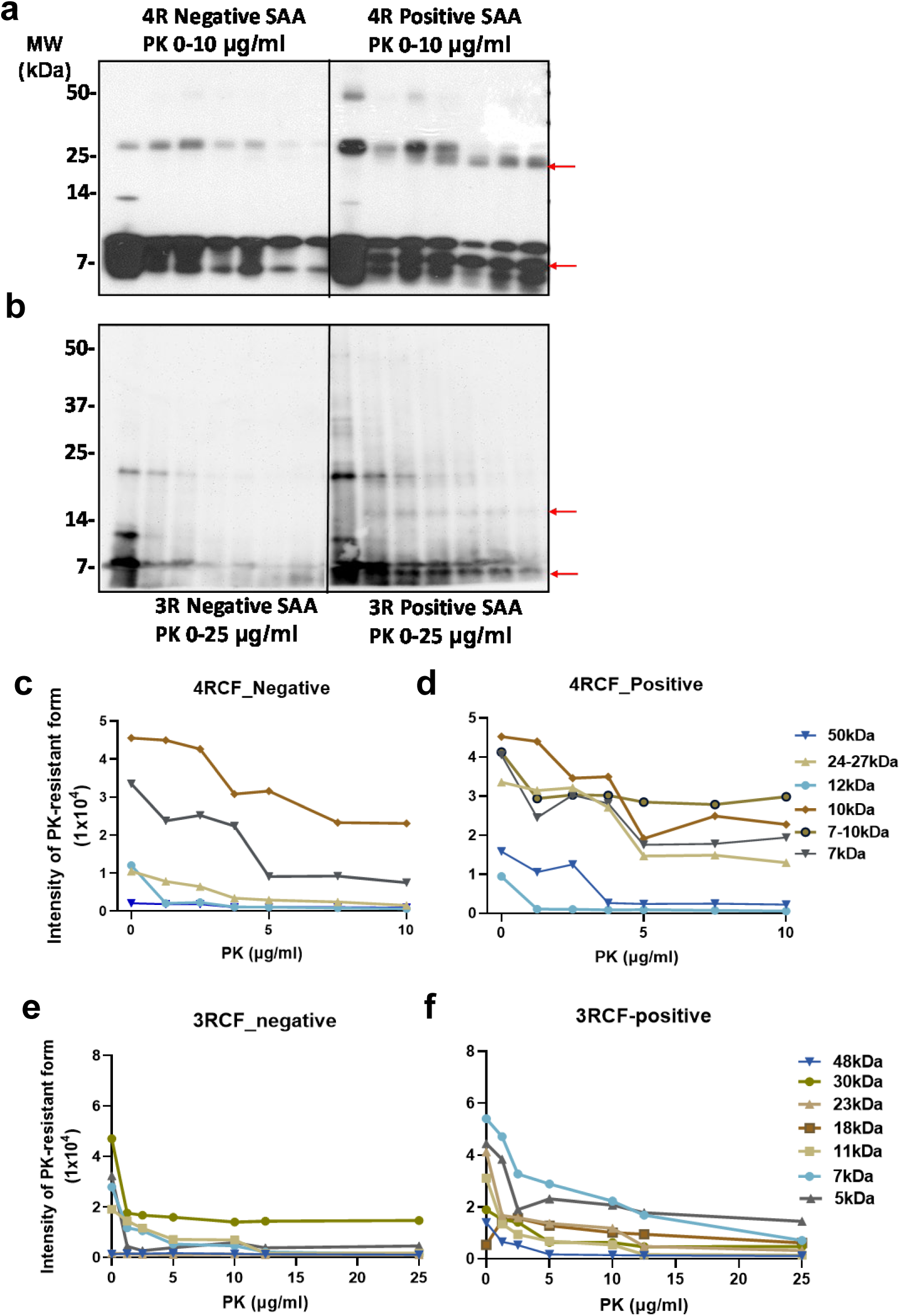
PK-resistance of the end products of 4RCF-/3RCF-based tau RT-QuIC assay of skin samples from tauopathies. **a** and **b** Western blotting of PK titration of 4RCF- (**a**) and 3RCF (**b**) - based RT-QuIC end product of skin samples from AD and non-AD controls. Probed with R3 and R4 antibodies against 3R or 4R tau isoforms, respectively. **c**-**f** Quantitative analysis of intensity of PK-resistant tau fragments at different molecular weights for 4R (**c**, **d**) and 3R (**e**, **f**) substrates.

Upon PK-treatment, for the non-AD end products, the intensity of the two lower monomeric tau bands was significantly decreased while they were still detected at PK of 10 μg/mL (Fig. 8a, c). The intensity of the tau band migrating at ~25-27 kDa was increased first up to PK of 2.5 μg/mL and then decreased, until became undetectable at PK of 10 μg/mL. The band migrating at ~12 kDa seemed to be completely PK-sensitive and no band was detectable even at the lowest PK concentration at 1.25 μg/mL. In contrast, after PK-treatment the end product of the RT-QuIC with AD skin samples showed the decreased intensity of the tau band migrating at ~25-27 kDa but generated additional smaller band migrating at about 24 kDa (Fig 8a, d, red arrow). This band is most likely derived from the truncation of the 25-27 kDa band since it was generated and increased over the increase in the PK concentration. In contrast with the non-AD end products, AD end product also exhibited an additional band between 7 kDa and 10 kDa bands migrating at about 8 kDa in the PK-treated AD skin end products (marked with the red arrow in Fig 8a). The intensity of this band was similar to that of 7 kDa and 10 kDa bands and showed no changes upon the increase in the PK concentration (Fig. 8a, d).

Regarding the end product of sTau-SAA using 3RCF as the substrate, without PK-treatment, the AD and non-AD samples all mainly exhibited 3 bands migrating at about 7 kDa, 10-11 kDa, and 22-23 kDa on the gel (Fig. 8b). According to the sequence of the 3RCF molecule, the molecular weight of the monomeric 3RCF should be 10.5 kDa. Therefore, the 7 kDa band could be a truncated fragment of 3FCF while 22-23 kDa band could be a dimer of 3RCF. Since we got high intensity of the low molecular weight bands (~7 kDa), we increased the PK concentration to 25 μg/mL for 3R and decreased the loading amounts of samples (Fig. 8b, e). The intensity of the 3 bands from non-AD end products all decreased while there was a faint band emerging, migrating at approximately 5 kDa over the increase in PK concentrations. The intensity of the 3 bands from AD skin tau RT-QuIC end products was also all decreased while there were two additional bands emerging, migrating at approximately 16-18 kDa and 5-6 kDa over the increase in PK concentrations (Fig. 8b, f, red arrows). The monomers of truncated 4RCF (at ~12 kDa) and 3RCF (at ~10-11 kDa) exhibited no resistance to PK treatment. However, the dimers at higher molecular weights demonstrate increased resistance to treatment, with the exception of the 4R negative end product dimers, which were digested at PK concentrations exceeding 7.5 µg/mL. Additionally, the low molecular weight bands below the monomers were more resistant to being digested within the chosen PK concentration range, especially with 4RCF.

### Conformational-stability assay of skin tau aggregates amplified by tau-SAA

We treated 4RCF (Fig. 9a) and 3RCF (Fig. 9b) tau-SAA end products with GdnHCl ranging from 0 to 3.2 M, followed by PK digestion at 10 µg/mL and quantitative analyses of GdnHCl/PK-resistant protein intensity of each treated sample. This approach is grounded on the principle that subtle differences in protein structure can be ascertained by assessing conformational stability when the protein is exposed to a denaturant such as GdnHCl at appropriate concentration ranges [23]. In the absence of GdnHCl and PK, 3 tau bands migrating at 48 kDa, 25 kDa and 7-12 kDa were observed for 4RCF-based RT-QuIC end products, while 2 bands migrating at 22-23 kDa and 5-10 kDa were detected in 3RCF-based RT-QuIC end products. In contrast, both AD skin 4RCF-/3RCF-based tau RT-QuIC end products were found to have multiple or smear bands above 30 kDa (Fig. 9a, b). After GdnHCl and PK-treatment, there were virtually no PK-resistant tau bands detectable from both non-AD 4RCF/3RCF-based RT-QuIC end products. In contrast, positive skin tau RT-QuIC from AD patients with either 4RCF or 3RCF as the substrate showed PK-resistant tau fragments, especially for bands migrating at 25 kDa or lower for low concentration of GdnHCl (Fig. 9). Notably, there was an additional partially PK-resistant tau fragment migrating between 7 kDa and 5 kDa bands for 3RCF-based RT-QuIC end products, which was not detectable in the 4RCF-based skin tau RT-QuIC end products (Fig. 9). The GdnHCl concentrations required to make half of the tau end product sensitive to PK, referred to as GdnHCl_1/2_, were 2.6 M for positive 4R tau and 2.3 M for positive 3R tau for bands migrating at 22-25 kDa and 10 kDa, indicating that the positive 4R tau end product is approximately 1.13-fold more stable than the 3R tau end product. But, the 3RCF-based RT-QuIC end products from AD cases generated stable 7 kDa band (Fig. 9b, f).

**Fig. 9.**
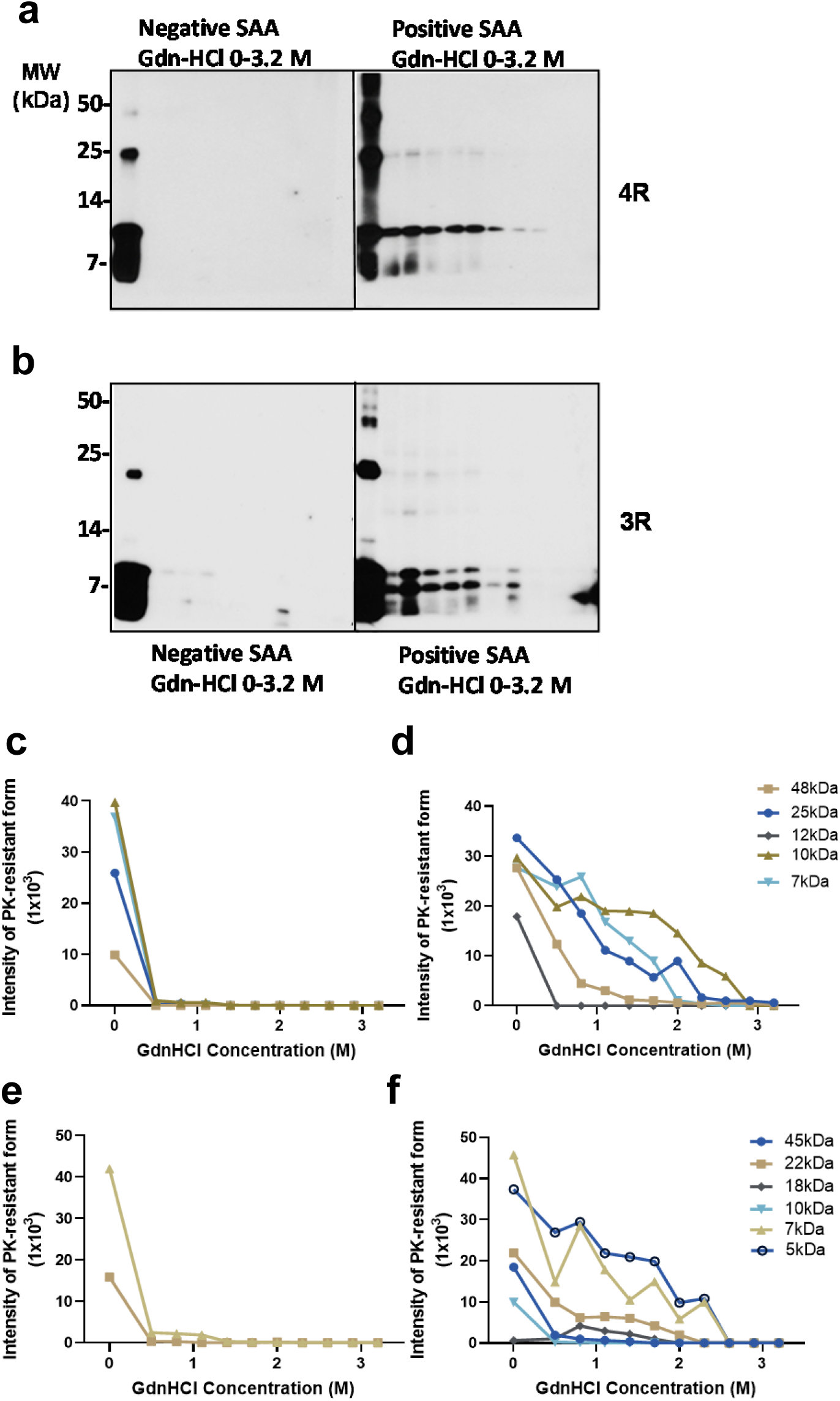
Conformational stability assay of 4RCF- or 3RCF-based RT-QuIC end products. **a** and **b** The 4RCF-(**a**)/3RCF(**b**)-based RT-QuIC end products were treated with different concentrations of GdnHCl and followed by PK digestion prior to Western blotting probed with anti-tau antibodies RD4 (**a**) and RD3 (**b**). **c**-**e** Quantitative analysis of Gdn/PK-resistant protein band intensity at different molecular weights for blots in **a** probed with antibodies against tau including RD4 (**c**, **d**) and RD3 (**e**, **f**).

### Fractionation of sTau-SAA end product by sucrose step gradient sedimentation

We further investigated the sedimentation of 4RCF and 3RCF skin tau SAA end products in sucrose step gradients, followed by western blotting of tau fragments in the sucrose step gradient fractions. The representative western blotting of sucrose gradient fractions from 4RCF-based RT-QuIC end products showed that significantly increased amounts of tau aggregates were detected in the bottom fraction 12 of AD and PSP samples but not in samples from other tauopathies (Fig. S4). In contrast, increased amounts of tau aggregates in the bottom fractions were mainly detected in the 3RCF-based RT-QuIC end products of skin samples from PiD and PSP but not in AD samples (Fig. S4).

## Discussion

There has been an unmet need to develop early and accurate biomarkers for use in AD clinical practice. This need goes beyond providing the information for diagnosis and prognosis of the disease, also for offering patients opportunities of clinical therapeutic trials and monitoring the therapeutic trial efficacies [20]. Following the recent encouraging Clarity AD trial report [13] and being at a crucial moment for developing new therapeutic strategies to prevent and delay AD progression [3], the need is becoming greater and more urgent.

Indeed, the recently developed AD-specific biomarkers used in clinical research greatly helped establish high certainty for diagnosing and monitoring AD pathology in living patients. Moreover, these biomarkers have also increased our understanding of the trajectory of accumulation and propagation of the misfolded Aβ and tau aggregates in AD [24]. Of the 3 broad categories of biomarkers in both 2018 initial and 2023 updated A/T/N research frameworks by the National Institute on Aging and the Alzheimer’s Association (NIA-AA), the detection of the misfolded proteins including Aβ and tau through the brain molecular imaging and body fluid examination was at the core of the framework [22, 60]. A new study evaluating the importance of various brain imaging modalities in monitoring cognition and predicting cognitive decline in memory clinics has observed that neocortical tau pathology is the main determinant of cognitive decline over time and the prognostic value of tau-PET surpassed all other neuroimaging measures [7]. One of the significant advances in the updated 2023 research framework is the addition of plasma biomarkers developed recently to the new version, while the 2018 version mainly depended on biomarkers from CSF and brain imaging [22, 60]. However, in addition to those disadvantages with brain imaging, CSF and Simoa-based plasma biomarkers as mentioned above, there are several other limitations as indicated in the 2023 update [60]. For instance, CT-PET is unable to detect neuritic plaques, low density and/or distributions of AD pathology restricted to medial temporal structures. Also, tau-PET cannot reliably differentiate among neuropathologically defined Braak stages I-III. Moreover, the plasma biomarkers cannot consistently discriminate among Braak stages I-IV in cognitively unimpaired subjects. It is also unclear whether the current biomarkers are available for all relevant neuropathologies and whether they reflect all respects of pathogenesis [60]. Lastly, the current list of the AD biomarkers in the updated framework still lacks any biomarkers that are able to reflect the pathological functions of misfolded proteins such as seeding activity of the pathogenic proteins either tau or Aβ, as the α-synuclein seed-amplification assay (SAA) for PD. As a result, there is a continuous search for new minimally or non-invasive biomarkers that are able to directly reflect the unique pathogenic features of the neurotoxic misfolded proteins themselves.

Using ultrasensitive RT-QuIC and/or PMCA, our previous studies have specifically detected minuscule amounts of misfolded proteins in the skin samples of other neurodegenerative diseases including prion disease and PD by detecting their seeding activity, a prion-like feature of misfolded proteins. For instance, we have demonstrated that the seeding activity of infectious prion protein (PrP^Sc^) and pathogenic α-synuclein (αSyn) can be detected in patients with Creutzfeldt-Jakob disease (CJD), prion-infected rodents and patients with PD or other synucleinopathies, respectively [5, 11, 26, 27, 39, 49, 50]. These findings have been verified by other groups [9, 30, 31, 54]. Inspired by these findings, we extended our study to the most common neurodegenerative disease AD and other tauopathies.

Our present study has made the following new findings. First, the levels of phosphorylated tau proteins detectable by western blotting in the autopsy skin tissue are significantly higher in AD than in non-AD tauopathies and normal controls; moreover, the gel profile of the detected phosphorylated tau is different between AD and controls including other non-AD tauopathies and normal subjects. Second, similar to tau in AD brains [25, 46, 53], autopsy skin tau-SA detected by RT-QuIC is significantly higher in tauopathies than in normal controls, suggesting sTau-SA can be used as a novel diagnostic biomarker for tauopathies. Third, sTau-SA can be detected in cases with PD and DLB but not with MSA although their seeding activity was significantly lower than that of AD while higher than normal controls. Fourth, the 3RCF substrate can be seeded by autopsy skin tau aggregates from all tauopathies, whereas the 4RCF substrate can be seeded by skin tau aggregates from AD, PSP and CBD but not from PiD. Fifth, the levels of tau-SA in the biopsy skin tissues in AD and PSP are significantly higher than those in the normal controls, implying the possibility of sTau-SA serving as a diagnostic biomarker for living tauopathy patients. Sixth, analysis of RT-QuIC end product reveals that sTau-SAA from AD forms tau oligomers and aggregates that are confirmed by our FTA, PK-treatment, conformational-stability assay, TEM, and sucrose gradient sedimentation. The ThT fluorescence intensity at the endpoint of the reaction correlates well with the tau aggregate dot intensity by FTA. Finally, our PK-treatment of skin tau RT-QuIC end products also exhibits greater amounts of PK-resistant tau fragments in AD than in normal controls. Moreover, conformational-stability assay reveals that skin tau can seed different strains in 4RCF and 3RCF substrates, respectively. Our findings raise several issues and implications as to the role of skin tau in the diagnosis and pathogenesis of tauopathies.

It has been shown that tau gene expression is significantly increased in the skin of males with aging [29]. Pathological tau deposits are detectable in peripheral organs including the aorta, liver, spleen, and stomach of AD patients but not in controls [35]. A tau isoform termed big tau, different from that in the brain was observed in the peripheral tissues of rodents [8, 16]. Phosphorylated tau has been observed in the skin of both AD patients and normal controls by immunohistochemistry, western blotting, and/or MALDI-MSI previously [1, 12, 43, 48]. Most of the above studies revealed that the skin tau gel profile is different from that of the brain tissues. Moreover, the gel profiles of skin tau from AD by western blotting were inconsistent among these studies [12, 43, 48]. For instance, Rodriguez-Leyva et al. reported two tau bands from AD migrating at ~55 kDa by AT8 and ~75 kDa by PHF antibodies with western blotting [43], whereas Dugger et al. observed a single band migrating at ~60 kDa with HT7, PHF-1, and pT231 antibodies and additional band at about 45 kDa with pT231 [12]. A recent study detected two major tau bands in the skin of AD and non-AD controls including 55 kDa and 70 kDa by western blotting with the following antibodies: Tau, Tau13, pT212, pS262 and pS404 for both 55 and 70 kDa and pS396 mainly for 70 kDa [48]. They all did not detect big-tau, the high molecular weight isoform of tau identified in peripheral nerves [15, 16, 48]. Our western blotting observed a skin tau gel profile with multiple bands in AD and other non-AD tauopathies, which was different from that of the previous observations [12, 43, 48]. The discrepancy among these studies could be due to different antibodies, experimental conditions, and sample processing used.

In addition to the bands observed in the previous studies including 70 kDa, 55 kDa and 45 kDa, we also demonstrated several other bands migrating at ~110 kDa, ~25 kDa, and even a high molecular weight band (HMWB) migrating above 110 kDa, respectively. The 110 kDa band could be equivalent to the so called big-tau, in terms of its molecular weight, which were predominately detected in all cases examined, even in the normal controls, except for a case from PiD. Notably, the HMWB was detected in cases from all groups except for the AD group examined. The other distinguishing feature of skin tau of AD patients on western blots was the gel profile of tau bands detected by the two anti-tau antibodies. It showed significantly increased levels of all tau bands in AD compared to that in other tauopathies and normal controls. The rich information about the levels and patterns of the tau molecules we identified in the skin tissues suggests that the skin is a useful specimen not only for investigating the role of tau in the pathogenesis of different tauopathies but also for developing differential diagnosis of AD from other tauopathies and controls.

Our current study is the first to show that tau from both autopsy and biopsy skin samples of AD and other non-AD tauopathies has a significantly higher seeding activity compared to normal controls, which suggests that the sTau-SA could be a novel diagnostic biomarker for tauopathies. Indeed, our 4RCF- or 3RCF-based RT-QuIC assay of the autopsy skin samples from AD and non-AD tauopathies yielded a sensitivity of 75-80% and a specificity of 95-100%, respectively. Tau-SA was also significantly higher in biopsy skin samples from tauopathies than from controls. Moreover, the skin tau-based RT-QuIC assay also can be used for the detection of comorbidity of AD and PD since we were able to detect skin tau-(this study) or αSyn-SA, depending on the substrate to be used [9, 11, 21, 26, 27, 30, 33, 49]. In other words, if the brain of a case has both tau- and αSyn-pathology, it is most likely that RT-QuIC will be able to detect both tau- and αSyn-SA in the skin of this case, which will indicate that the case examined has comorbidity of AD and PD. However, before we can make the comorbidity diagnosis reliable, we need to make sure that it does not result from cross-seeding between tau and αSyn. This is because cross-seeding between the two will let tau-seeds trigger αSyn aggregation, and vice versa. In addition, it is worth noting that while skin tau from PiD mainly containing 3R-dominated tau aggregates in the brain was able to seed 3RCF but not 4RCF substrate by RT-QuIC, skin tau from AD mainly with a mixture of 3R/4R tau and from PSP/CBD characterized by the 4R-dominated tau aggregates in the brain was able to seed not only 4RCF but also 3RCF substrate.

Our analysis of skin-tau RT-QuIC end products confirmed that the increased tau-SA by tau-SAA was companied with the formation of tau aggregates as shown by our FTA, PK-treatment, sucrose gradient fractionation, and conformational-stability assays. Our TEM did not reveal protofibrils or mature fibrils in 3RCF- or 4RCF-based RT-QuIC end products of skin tau from AD cases although the end products of our RT-QuIC of skin αSyn from PD cases as a control showed protofibrils and mature fibrils. They seemed to form soluble oligomer-like structures, as shown in the literature [2, 6]. Our previous study showed that recombinant 4RCF fragments can spontaneously form mature fibrils *in vitro* in about 2 weeks [53]. Based on PK-digestion of skin tau RT-QuIC end products, positive RT-QuIC end products generated additional PK-resistant tau fragments. Moreover, the 3RCF-and 4RCF-based RT-QuIC end products generated different PK-resistant fragments. It will be interesting to determine whether end product of tau-SAA with brain and skin samples would generate the same or different structural and physicochemical features in the future.

In conclusion, our study underscores the potential of skin samples as a minimally invasive and accessible source for the diagnosis of tauopathies. The identification of phosphorylated tau and tau seeding activity in skin samples could facilitate the development of innovative diagnostic approaches for tauopathies. Further research with larger patient cohorts and optimization of detection methods will be critical in establishing the clinical utility of a skin-based approach for diagnosing and differentiating between different tauopathies.

## Acknowledgments

We are grateful to all donors and their families for their skin tissue donations, and to Drs. Thomas G. Beach and Geidy E. Serrano from Banner Sun Health Research Institute, Sun City, AZ, USA for providing autopsy scalp skin tissues from cadavers with tauopathies, synucleinopathies, and controls as well as related information.

## Author Contributions

W.Q.Z. conceived and designed the study. W.Z. and W.Q.Z. developed and interpreted RT-QuIC analysis of the skin samples. W.Z. performed RT-QuIC analysis of the skin samples. W.Z., M.G., T.G., S.Z.A.S. and W.Q.Z developed, performed, and interpreted the analyses of RT-QuIC end products of the skin samples. L.W. and B.X. designed, developed and performed preparation of human recombinant 4RCF and 3RCF proteins and identified site-specific phospho-tau epitopes. S.G., R.L. and V.D. provided biopsy skin samples and related clinical data. W.Z. and W.Q.Z. wrote the first version of the paper. All authors critically reviewed, revised, and approved the final version of the manuscript.

## Data and materials availability

All materials used in this study will be made available subject to a materials transfer agreement.

