## Supplemental information for "Seeding Activity of Skin Misfolded Tau as a Novel Biomarker for Tauopathies"

**This file includes:**

Supplementary Text

Figs. S1 to S4

**Figure S1**. **Western blotting of tau in the skin of tauopathies without primary antibodies.** ***a***: Conventional western blotting of skin homogenates probed by only secondary antibody sheep anti-mouse IgG. ***b***: Conventional western blotting of skin homogenates probed by only secondary antibody donkey anti-rabbit IgG. The exposure time of the blots were 90 min to make sure that the protein bands would be visible if present.

**Figure S2. Assessment of the full-length and truncated tau as the substrate of skin tau RT-QuIC assay.** ***a***: Endpoint tau-SAA results for skin tissues using truncated 4RCF/3RCF tau fragments and full-length 2N3R/2N4R tau proteins, as indicated by tau ThT fluorescence measurements at the endpoint of the RT-QuIC assay with AD skin tissues. ***b***: Tau-SAA lag phase of autopsied AD skin tissues utilizing truncated 4RCF/3RCF tau fragments and full-length 2N3R/2N4R tau proteins. The lag phase was determined from the kinetic ThT fluorescence curves started to increase through the RT-QuIC assay. ***: $p < 0.004$

**Figure S3: Transmission electron microscopy (TEM) of various SAA end products.** a and b: Representative image of positive 3RCF (***a***) and 4RCF (***b***) SAA end products, displaying oligomer-like structures. ***c***: Representative image of negative αSyn SAA end product that shows no oligomers or fibrils. ***d***: Representative image of positive αSyn SAA end product, highlighting the existence of mature fibrils. Scale bar: 200 nm.

**Figure S4. Western blotting of sucrose gradient sedimentation fractions of skin tau RT-QuIC end-products from different tauopathies.** ***a***: Representative western blotting of sucrose gradient fractionation of 4RCF-based RT-QuIC end-products from different tauopathies including AD, PSP, PiD, CBD and NC. ***b***: Densitometric analysis of tau bands from 4R SAA end-product distribution across different sucrose gradient fractions from different tauopathies including AD, PSP, PiD, CBD and NC, 3 for each. ***c***: Representative western blotting of sucrose gradient fractionation of 3RCF-based RT-QuIC end-products from different tauopathies including AD, PSP, PiD, CBD and NC. ***d***: Densitometric analysis of tau bands from 3R SAA end-product distribution across different sucrose gradient fractions from different tauopathies including AD, PSP, PiD, CBD and NC, 3 for each.

SI

Figure S1

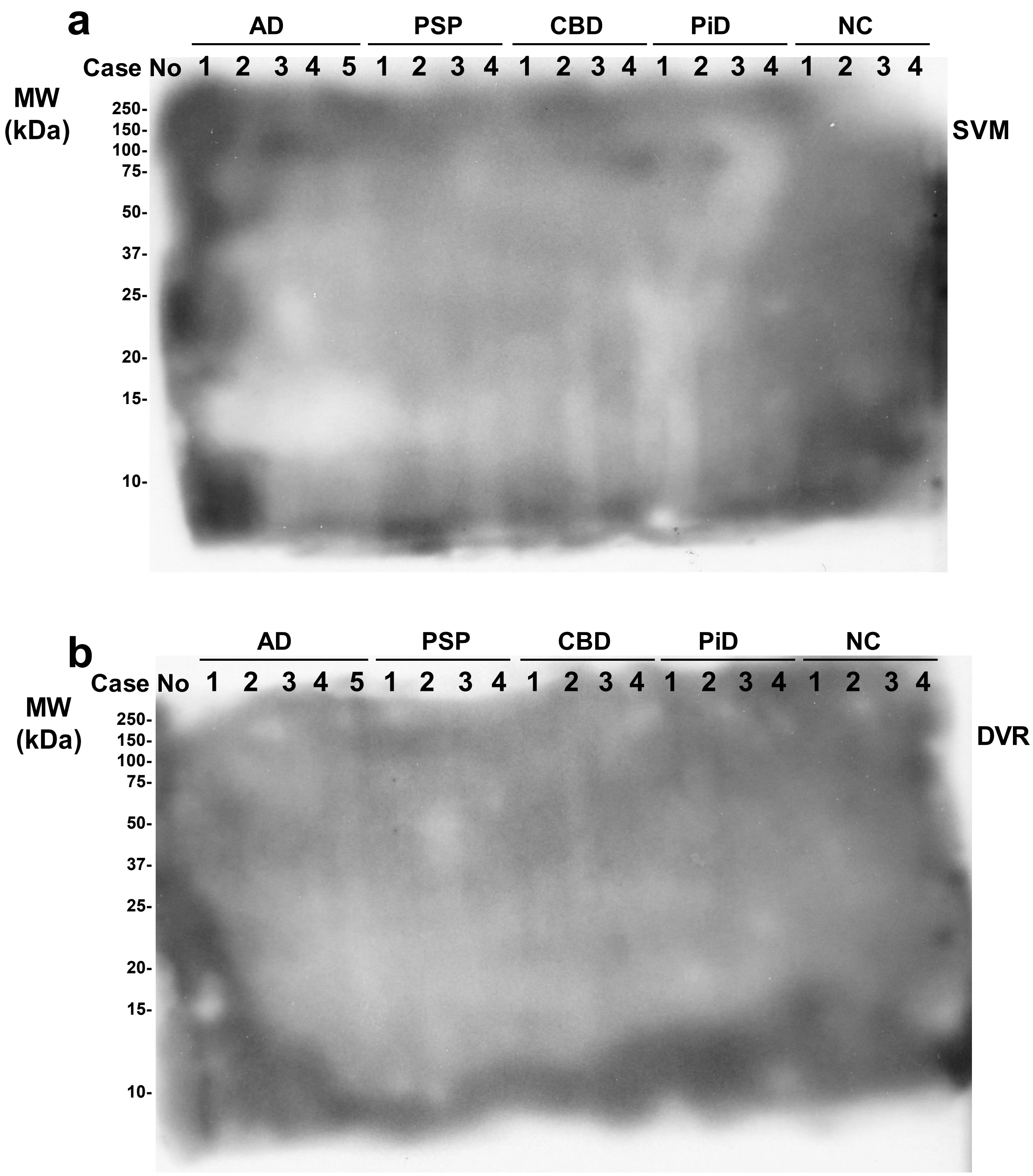
a
AD
PSP
CBD
PiD
NC
Case No 1 2 3 4 5 1 2 3 4 1 2 3 4 1 2 3 4 1 2 3 4
MW (kDa)
250-
150-
100-
75-
50-
37-
25-
20-
15-
10-
SVM
b
AD
PSP
CBD
PiD
NC
Case No 1 2 3 4 5 1 2 3 4 1 2 3 4 1 2 3 4 1 2 3 4
MW (kDa)
250-
150-
100-
75-
50-
37-
25-
20-
15-
10-
DVR

Figure S2

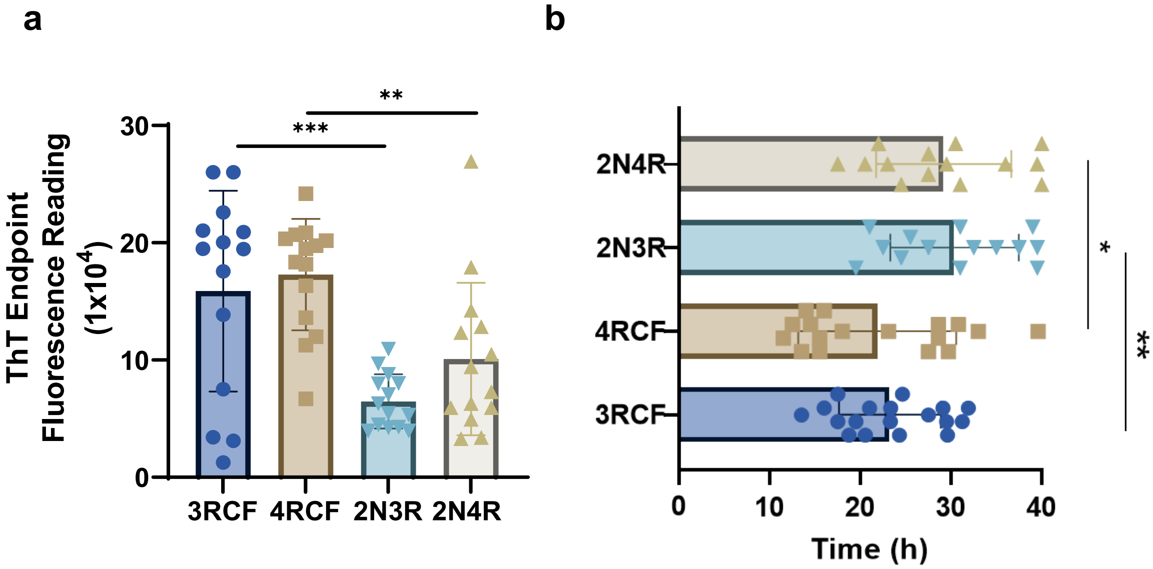
a
ThT Endpoint Fluorescence Reading (1x10^4)
30
20
10
0
***
**
3RCF
4RCF
2N3R
2N4R
b
2N4R
2N3R
4RCF
3RCF
*
**
0
10
20
30
40
Time (h)

Figure S3

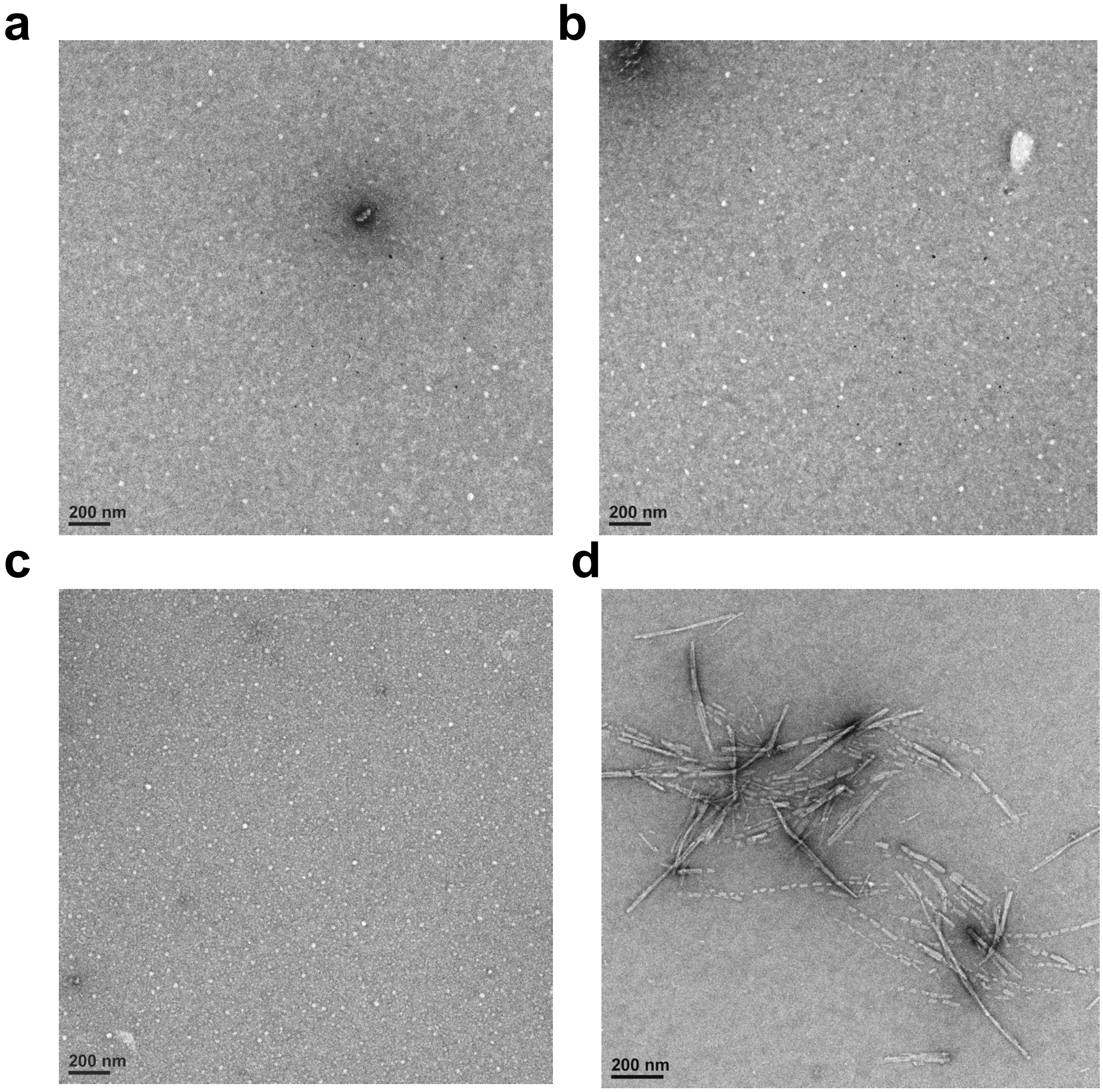

Figure S4

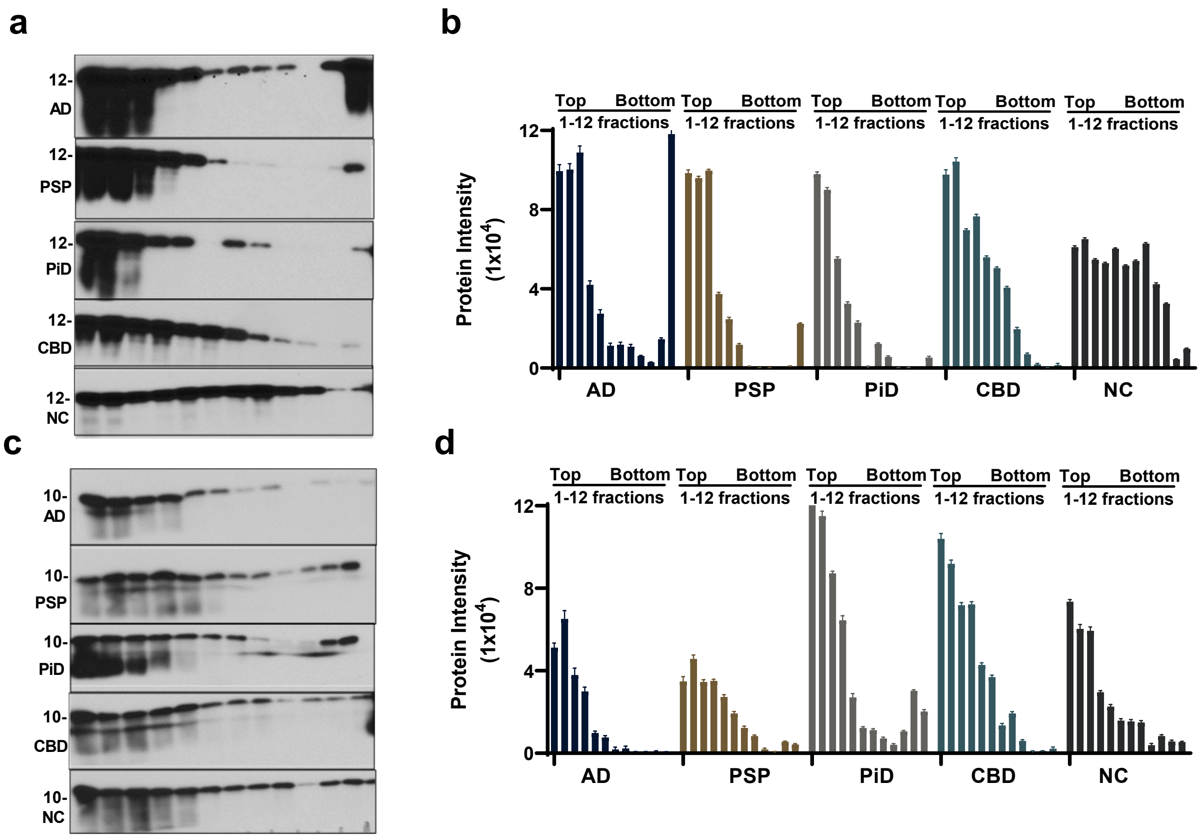

a
12-
AD
12-
PSP
12-
PiD
12-
CBD
12-
NC
b
Top Bottom
1-12 fractions
Protein Intensity (1x10^4)
12
8
4
0
AD
PSP
PiD
CBD
NC
c
10-
AD
10-
PSP
10-
PiD
10-
CBD
10-
NC
d
Top Bottom
1-12 fractions
Protein Intensity (1x10^4)
12
8
4
0
AD
PSP
PiD
CBD
NC
